# Overexpression of the violaxanthin de-epoxidase confers faster NPQ and photosynthesis induction in rice

**DOI:** 10.1101/2023.04.26.538496

**Authors:** Duanfeng Xin, Faming Chen, Pengfei Zhou, Zai Shi, Qiming Tang, Xin-Guang Zhu

## Abstract

Plants in the field experience fluctuations in light conditions. Plants with swift responses to dynamic light conditions usually gain competitive advantage in the field. The dynamic photosynthesis can be reflected in dynamic changes in almost all processes in photosynthesis, with dominant factors being dynamic changes in non-photochemical quenching, Rubisco activation and stomatal dynamics. Each of these dynamic responses is controlled by defined mechanisms. In this study, we showed that overexpression of VDE, an enzyme involved in the xanthophyll cycle and non-photochemical quenching, in rice resulted in faster NPQ induction, less photoinhibition, and faster photosynthesis induction, which together caused increased biomass production by about 11∼16%. This study, demonstrating a previously unnoticed role of VDE in altering the dynamics of photosynthesis induction besides decreasing photoinhibition, offers a potential strategy to improve canopy photosynthesis through capitalizing the ability of plants to use dynamic light.

**Highlight:** The overexpression of VDE in rice resulted in faster NPQ induction, less photoinhibition, and faster photosynthesis induction, which together caused increased biomass in field.

## Introduction

Field grown plants experience constantly fluctuating light as a result of the cloud, wind and so on [1, 2]. Leaves constantly adjust photosynthetic machineries to ambient light levels through different mechanisms, such as altering chloroplast positions, photosynthetic antenna sizes, activation status of different enzymes related to photosynthesis etc. The challenge is that the adjustment is not instantaneous. For example, when leaves move into full light from shade, the carbon assimilation rate does not reach its maximum value immediately, but rises to a stable state in several minutes [3, 4]. Similarly, when the leaves go from full sunlight to shade, the photosystem II operation efficiency drops to a lower level and it takes a few minutes to relax non-photochemical quenching and hence regain higher photosynthetic efficiency [1, 5]. The slow response to frequent light transition may reduce the potential carbon assimilation in wheat, cowpea, African cassava and others by 13∼32% [6-14].

There are several factors involved in responses of photosynthesis to dynamic light, which include metabolic processes, gas diffusion, NPQ and others [5]. Metabolic processes include the activation of Calvin-Benson cycle (CBB cycle) and electron transport processes [3, 6, 15-19]. The activation of CBB cycle involves the dynamic regulations of different enzymes, especially the ribulose 1,5-bisphosphate carboxylase/oxygenase (Rubisco), whose activity is heavily influenced by the activity of Rubisco activase (RCA)[20]. The enhancement of RCA capacity could lead to faster photosynthesis induction in rice and tobacco [3, 17, 21]. Rubisco activation process is not only influenced by RCA, but also the CO_2_ level around Rubisco, which further is influenced by the diffusion of gas from air to the site of CO_2_ fixation by Rubisco. The gas diffusion is usually influenced by two factors: mesophyll conductance (g_m_) and stomata conductance (g_s_) [22, 23]. In Arabidopsis and tobacco, g_m_ is not a limiting factor during photosynthesis induction while g_s_ and electron transport rate are [15]. In African cassava, g_s_ was again a main limitation under fluctuating light, which is in contrast to the steady state conditions, where Rubisco and g_m_ are the main limitations for photosynthesis [10]. Consistent with this, improved stomatal opening enhanced the photosynthesis induction and biomass production under natural fluctuating light in both Arabidopsis [23] and rice [16].

The dynamic change of non-photochemical quenching (NPQ) is another major factor influencing photosynthetic performance under dynamic light [13, 24-27]. Leaves under full sunlight receive more solar energy than they can use, resulting in a large proportion of incident light dissipated in the form of heat, which help decreases the production of oxidizing radicals, chemicals damaging photosynthetic apparatus [28-31]. NPQ is regulated by pigment composition of thylakoid membrane, the amount of PsbS, and the proton gradient across the thylakoid membrane [29, 32-35]. When leaves are exposed to high light, proton concentration gradient across the thylakoid membrane increases to activate the xanthophyll cycle, resulting in conversion of violaxanthin (Vio) into zeaxanthin (Zea) by violaxanthin de-epoxidase (VDE) and increased NPQ [34, 36, 37]. In turn, Zea can be converted back to Vio by Zeaxanthin epoxidase (ZEP), with NADPH and oxygen as cofactors [38]. The antheraxanthin (Ant) is the intermediate metabolite during the conversion between Vio and Zea[34]. In addition, high concentrations of protons can bind and protonate PsbS proteins to rebuild the microstructures of the super complex of photosystem II (PSII) and light harvested complex II (LHCII) [39-42]. The protonation of PsbS and the de-epoxidation of Vio form energy-dependent quenching (qE), the fastest part of NPQ [28, 31, 37]. Besides qE, Zea-dependent quenching (qZ) and photoinhibition-dependent quenching (qI) are also the vital parts of NPQ [37, 43]. Under the normal growth of a plant, NPQ and the repairing process need to be balanced to avoid photoinhibition and correspondingly decreased photosynthesis [44, 45]. When plants experience a sudden high light, the balance is disrupted and photoinhibition occurs [44, 46, 47]. Photoinhibition means here the rapid decline of PSII activity under stress [44, 48, 49]. Though NPQ and photoinhibition are independent from each other, photoinhibition can be affected when NPQ is genetically altered [24, 39, 40, 50]. For example, enhanced NPQ can increase productivity by alleviating photoinhibition[24, 27, 51, 52]. As expected here, the NPQ needs to be fine-tuned, and supera-optimal NPQ levels can also reduce the operating PSII quantum yield [24, 25, 27, 53, 54].

Given the importance of NPQ in controlling photosynthetic efficiency under dynamic light and the inherent temporal and spatial heterogeneity of light inside a canopy, manipulating NPQ dynamics is regarded as a major option to improve photosynthetic efficiency[1, 24, 25, 27]. The stacking of three NPQ-related proteins from *Arabidopsis thaliana*, VDE, ZEP and PsbS could accelerate the response of NPQ and photosynthesis under fluctuating light, resulting in an increase of biomass by about 15∼20% in tobacco in the field [25]. A recent study further showed that improving the dynamics of NPQ can improve soybean photosynthesis and yield in the field as well [27]. However, the accelerated NPQ by the three genes did not result in improved growth and biomass accumulation in Arabidopsis [53] and potato [54]. Similar to the effects of over-expressing these three genes, over-expression of some single genes involved in NPQ regulation also provides promising results on photosynthesis enhancement. The overexpression of PsbS in rice alone also resulted in higher NPQ with increasing canopy radiation use efficiency and yield under fluctuating light [24]. The overexpression of VDE from cucumber and *Lycium chinense* respectively improved the tolerance to high light and drought in Arabidopsis [55, 56]. Consistently, the *Cerasus humilis* derived VDE enhanced the tolerance to drought and salt stress in transgenic Arabidopsis [57]. Overexpressing a tomato VDE or a peanut VDE gene alleviated the sensitivity of PSII photoinhibition to stress in tobacco [51, 58]. The overexpression of tomato-derived VDE increased the photosynthetic efficiency under high light or chilling stress in tobacco [36, 51, 55, 57, 59, 60].

In this study, we showed that, in rice with overexpression of VDE, besides the well-known effect of increased NPQ initiation, there is an increased photosynthesis induction, which resulted in increased biomass production, suggesting that manipulation of VDE alone might be a potential approach to improve whole canopy photosynthetic CO_2_ uptake for greater yield.

## Materials and methods

### Plant material

We used rice (*O. sativa* L. cv. Xiushui134) in this study. For the overexpression vector, the coding region of VDE was amplified from the cDNA of rice by gene specific primers CDS-F/R and inserted into an OE vector containing maize ubiquitin (Ubi-1) promoter and NOS terminator, which was derived from pCAMBIA1300 and fused with a 3xFLAG tag using ClonExpress II One Step Cloning Kit (C112-01, Vazyme, China). The bacterial resistance of the vector was kanamycin (Kan) and the plant resistance was hygromycin (Hyg). Then the OE vector was transformed into rice via the Agrobacterium-mediated method[61].

The sequence of related primers:

> CDS-F: AGCTCGGTACCCGGGGATCCATGATGCCGCGGCAGTGCGG.
>
> CDS-R: AGGTCGACTCTAGAGGATCCCCTTAGTTTCCTTATTGGCA.

### Growth conditions in the field and greenhouse

In our field experiment, rice was grown at the Songjiang experimental station (30°56’ N, 121°8⍰ E) of the Chinese Academy of Sciences Center for Excellence for Molecular Plant Sciences, Shanghai, China. All seeds were sown on seedbeds after germination on May the 22^nd^ of 2021. Seedlings were transplanted on June 19^th^ of 2021. Each plot was designed in a 7×7 layout (49 plants) with the planting row distance and column distance being 20cm. WT, OE-VDE1, OE-VDE3 and OE-VDE5 were planted in sequence and repeated 5 times to avoid position effects.

In the greenhouse, the temperature was set to 27 °C and the relative humidity was around 60-70%. Light was turn on from 7 a.m. to 7 p.m. and the photosynthetic photon flux density at the top of the plants was about 400 μmol m^-2^ s^-1^. The rice seeds were geminated in petri dishes with distilled water at 30 °C under dark for 4 days and then transferred to ½MS solution. The ½MS solution was refreshed every week.

The plants used for the measurements of gas exchange and chlorophyll florescence were transplanted to 30cm*40cm square pots with 15 liters paddy soil mixed with 5g fertilizer (N:P_2_O_5_:K_2_O=20:20:20) as base fertilizer after growing in pots with ½MS solution for 1 month. Rice seedlings were grown in a 3×4 layout (3 biological repeats for each genotype, i.e., WT, OE-VDE-1, OE-VDE-3 and OE-VDE-5) in each pot. We used 4 pots. Plants were grown with different orders in different pots to avoid position effects.

For the measurement of NPQ under normal light, plants were grown in pots with ½MS solution for 50 days in greenhouse. For the measurement of NPQ under high light, plants grown in pots with ½MS solution for 45 days were moved to high light (1000 μmol m^-2^ s^-1^) for 3 days in greenhouse. For the measurement of NPQ in field, the leaf samples (12 cm, the middle part of the last fully expanded leaf) were taken from the field at about 7 a.m. and soaked in 0.1% X-Triton solution under dark before measurement.

### Chlorophyll fluorescence measurement by Pulse Amplitude Modulation (PAM)

We also measured the time course of chlorophyll fluorescence-based parameters in PSII by Dual-PAM-100 (Heinz Walz GmbH, Germany). We used newest fully expanded leaves at the tillering stages. Before the measurements, leaves were first dark adapted for 2 hours. The minimal fluorescence yields (Fo) were recorded by the measure light (630 nm, 24 μmol m^-2^ s^-1^) and the maximal fluorescence yields (Fm) was recorded during the first saturation pulse (630 nm, 10000 μmol m^-2^ s^-1^, 500 ms). The actinic light (630 nm, 1300 μmol m^-2^ s^-1^) was generated for 10 mins followed by 6-min darkness, during which the saturation pulse was launched every 20 s. The steady-state fluorescence yields (F) and the maximal fluorescence yields under light (Fm’) were recorded after each saturation pulse [62, 63].

### Gas exchange measurement

The newest fully expanded leaves at the tillering stage were selected for measurements. The leaf temperature was set at 27 °C with an ambient CO_2_ level of 410 μmol mol^-1^. LI-6800 (LI-6800 Portable Photosynthesis System, LI-COR, USA) with a 6800-01A Multiphase Flash Fluorometer and Chamber was used for the measurements. F and Fm’ were recorded after each saturation pulse (625 nm, 10000 μmol m^-2^ s^-1^, 250 kHz) under the actinic light (625 nm). Fo was recorded by the measure light (625 nm, 0.05 μmol m^-2^ s^-1^) and Fm was recorded after the saturation pulse.

For the measurement of photosynthetic CO_2_ uptake rate (*A*) versus photosynthetic photon flux density (Q) (*A*-Q) curves, the leaf was first given a 10-minute light adaption under a photosynthetic photon flux density of 2000 μmol m^-2^ s^-1^. Then, Fm’ and gas exchange parameters were recorded and light intensity was changed from 2000 to 1500, 1200, 1000, 800, 600, 400, 200, 150, 120, 90, 60, 30, 15 and 0 μmol m^-2^ s^-1^. Between measurements at each light levels, leaves were light adapted for about 75 s. Fo and Fm was recorded after dark adaptation for 2 h.

During the measurement of dynamic photosynthesis after leaves were switched from low to high light, the leaves were first light adapted under a PPFD of 100 μmol m s for 4 min; then the PPFD was switched to 2000 μmol m s for 10min. F and Fm’ were recorded every 12 s while gas exchange parameters were recorded every 3 s. Fo and Fm was recorded after dark-adaption for 2 h.

For the accurate measurement of 1-qL during the photosynthesis induction from low to high light, leaves were first light adapted for 4min under a PPFD of 100 μmol/ m^-2^ s^-1^. Then the PPFD was switched to 2000 μmol/ m^-2^ s^-1^ for 336 s. The gas exchange parameters were recorded every 3 s and the Fo’ were recorded via launching a far-red light (735 nm) at the 33^rd^ s and 336^th^ s. Fo and Fm was recorded after dark-adaption more than 2 h.

We calculated the chlorophyll fluorescence-based parameters based on the following formulas [63, 64]:

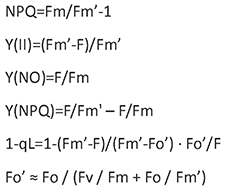

### Pigment analysis

Leaf discs each with an area of around 2 cm^2^ were collected with a leaf punch from the newest fully expanded leaves in the tillering stage in field and frozen in liquid nitrogen immediately. The frozen leaves were grounded into powder in liquid nitrogen. Then the samples were extracted with 1ml 80% ice-cold acetone and shaken slowly at 4℃ for 3h. After 5 min of centrifugation at 13,000 rpm, the supernatant was filtered with 0.22μm nylon strainer. Finally, the pigment samples were analyzed with reverse-phase high-performance liquid chromatography (HPLC, Agilent 1260, USA). A C30 column (92114-630) was used during the analysis. The mobile phase was composed of 100% methanol and ran for 10min at a flow rate of 1ml/min. Eluted pigments were detected at 440 nm. The protocol used was provided by Beijing PZD Science and Technology Co., Ltd.

### Western blotting of VDE

10 cm^2^ leave samples were collected from the newest fully expanded leaves at the tillering stage in the field and grounded into powder within liquid nitrogen. We then added 400 μL SDS protein extraction buffer (4% SDS, 50mM Tris-HCl pH 7.4, 2 mM EDTA, 100 mM NaCl, 10mM DTT) to lyse the samples. After fully mixed, the lysed samples were heated to 95℃ for 5 min. After that, the samples were centrifuged and the supernatant was removed into a new 1.5 ml tube. The protein quantification was performed via the Detergent Compatible Bradford Protein Assay Kit (P0006C, Beyotime, China) and Western blotting was performed according to the manufacturer’s protocol from ABclonal (https://abclonal.com/Uploads/protocol/WB_protocol.pdf). An anti-VDE antibody (AS15 3091, Agrisera, Sweden) and HRP Goat Anti-Rabbit IgG (H+L) (AS014, ABclonal, China) were used to examine the changes of protein levels in different OE-VDE lines and WT.

### Gene expression analysis of VDE

The leaf samples were collected from the newest fully expanded leaves at the tillering stage in the field and immediately frozen into liquid nitrogen. We used GeneJET Plant RNA Purification Kit (K0802, Thermo Scientific, USA) for RNA extraction following manufacturer’s protocol (https://www.thermofisher.cn/order/catalog/product/K0802). The HiScript III All-in-one RT SuperMix (R333-01, Vazyme, China) was used for reverse transcription and Taq Pro Universal SYBR qPCR Master Mix (Q712, Vazyme, China) was used for Reverse transcription quantitative PCR (RT-qPCR). OsActin1 (Os03g0718100) was used as control.

> The primers used for RT-qPCR:
>
> ACTIN1- qRT-F: TGAGTAACCACGCTCCGTCA.
>
> ACTIN1- qRT-R: CCTTCAACACCCCTGCTATG.
>
> VDE-qRT-F: CCGTGGCAGAAATGATGCATG.
>
> VDE-qRT-R: GTCGGTCCTGATGAACGTGG.

### Measurement of agronomic traits

At the harvest stage, 60 plants of each line were selected from 5 plots with 12 plants taken from each plot. First, the plant height and the panicle number of each plant were measured. Then, the aboveground biomass and grain were dried at 70 °C for 4 days and at 40 °C for 7 days until the samples were completely dried. Finally, the aboveground biomass and grain yield of each plant were measured.

For the measurement of specific leaf weight (SLW), leaf samples at the heading stage were collected and dried in oven at 110℃ for 1 h, and then at 70℃ for 4 days until the samples were completely dry. The dry samples were grounded into powder for the measurement of leaf nitrogen content (LNC) by a vario ISOTOPE CUBE elemental analyzer (elementar, Germany).

During all these measurements, we took samples from plants growing in the middle of plots to avoid boundary effects.

### Statistical analysis

Data were analyzed via the two-sided t-test in Microsoft Office Excel software. When *p* < 0.05, the difference was considered to be significant. The related figures were drawn by the GraphPad Prism software (version 8.0.2).

## Results

### Overexpression of VDE increased the content of Ant and Zea in rice leaves

We overexpressed VDE (Os04g0379700), homologous with AtVDE (AT1G08550) (Fig 1A and Fig. S1), and obtained three independent VDE-overexpression lines (OE-VDE- 1/3/5), in which their expression levels were significantly higher compared with those in Wild type (WT) (Fig 1B and 1C). We quantified the pigment contents of rice leaves using High Performance Liquid Chromatography (HPLC) (Fig. S6). As expected, in the OE-VDE lines, higher amount of Ant and Zea were shown, while less Vio was detected. The contents of Zea and Ant in OE-VDE lines increased by 260-410% and 100-130% compared to WT, respectively (Fig. 2A∼2C). In contrast, the contents of lutein (Lut) and chlorophyll did not show significant difference between OE-VDE and WT (Supplementary Fig. S2A∼2D). The xanthophyll cycle de-epoxidation state (DES)[34, 65] was increased by 200-270% in leaves of OE-VDE lines compared to that of WT (Fig. 2D). All these are consistent with the biological function of VDE, catalyzing the conversion from Vio to Ant and Zea (Fig. 2E).

**Fig 1.**
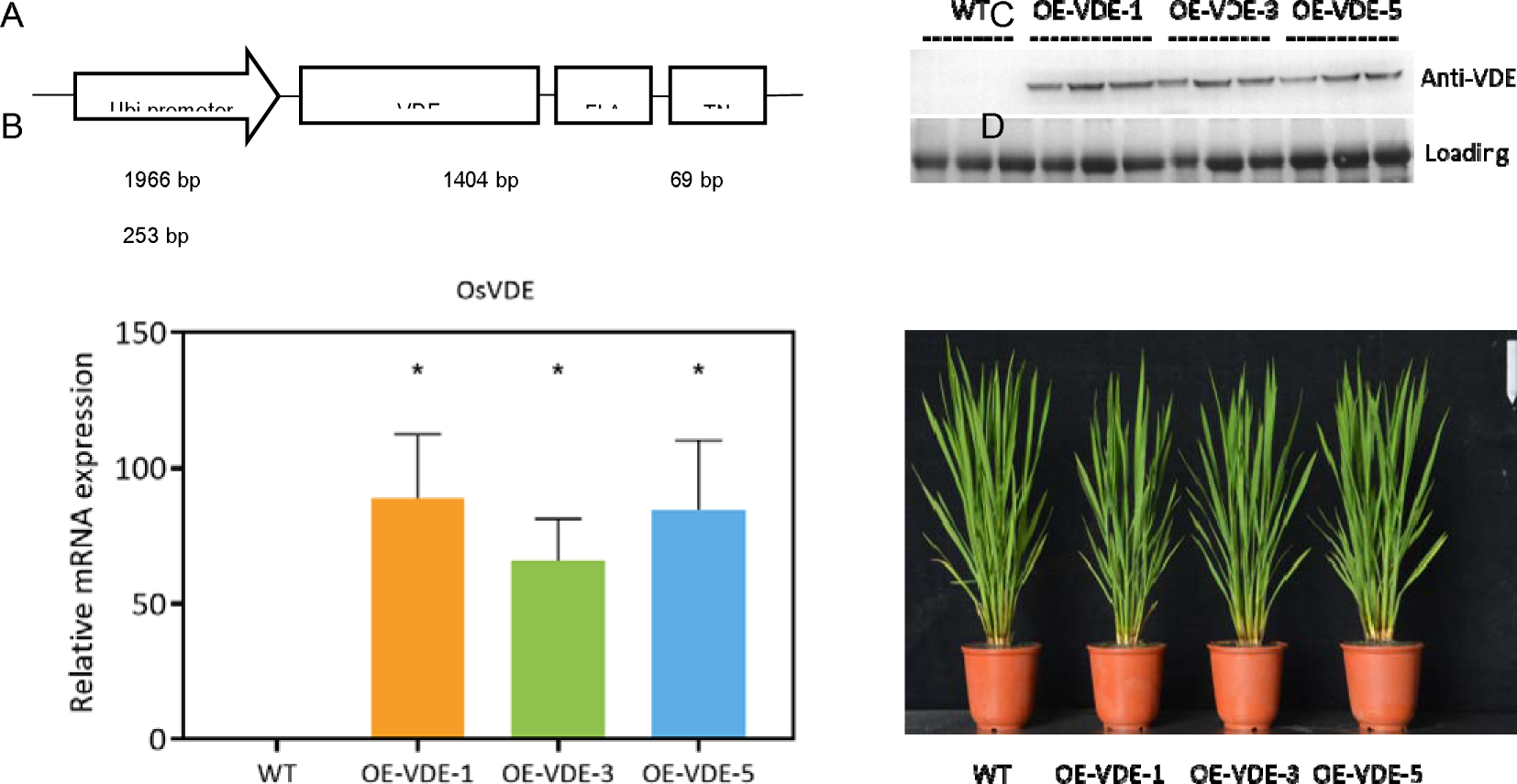
Construction and identification of Rice VDE overexpression transgenic line (OE-VDE-1, OE-VDE-3, OE-VDE-5). (A) The segment used to overexpress OsVDE which was integrated into pCAMBIA1300 vector; (B) The RNA quantification of the OE-VDE lines by qrt-PCR, n = 3, *: p<0.05 (Student’s t test); (C) Western blotting of OsVDE protein from the OE-VDE lines and WT by using the anti-VDE antibody. The protein samples were added equally and Rubisco staining shown below served as the loading control. The last expanded rice leaves at tillering stage in the field were collected at 2 p.m. on a sunny day. OE-VDE: the transgenic rice overexpressing VDE; WT: wild type. (D) Phenotype of OE-VDE lines. Picture was taken from typical plants of each line in field at 60 days after transplant (DAT), in Shanghai.

**Fig. 2.**
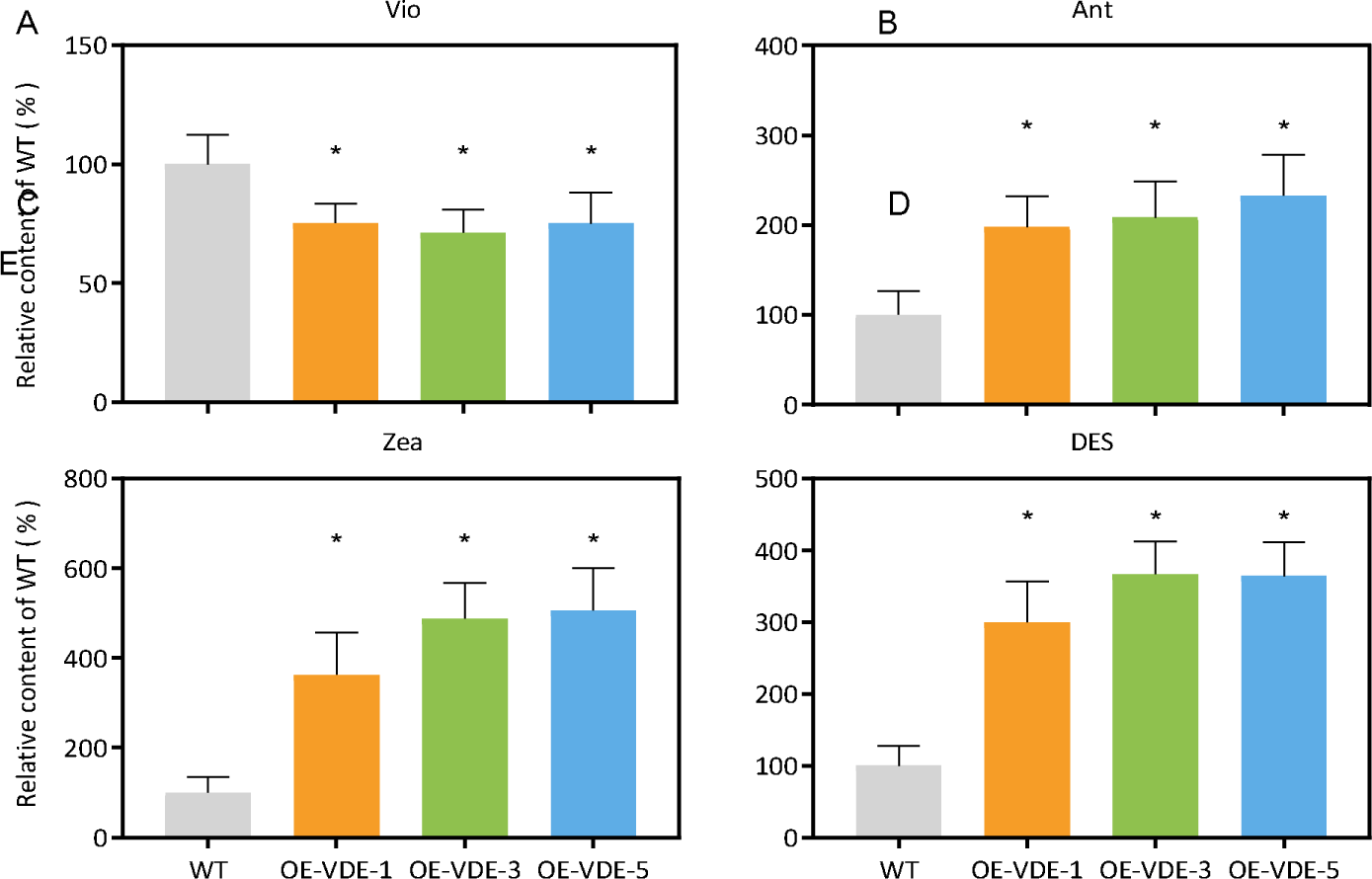
Change in xanthophyll cycle pigment content and DES in the OE-VDE lines compared with WT. (A) Vio, (B) Ant, (C) Zea, (D) DES (de-epoxidation state). The last expanded rice leaves at tillering stage in the field were collected at 2 p.m. on a sunny day. n = 6∼7, *: p<0.05 (Student’s t test). (E) The model of xanthophyll cycle.

### Overexpression of VDE accelerates NPQ initiation

We further examined whether the overexpression of VDE changed the response of NPQ in rice leaves, we measured the NPQ dynamic curves of rice plants grown in greenhouse. Result show that the speed of NPQ induction in the OE-VDE lines were faster than that of WT in the first 40^th^∼80^th^ s, but there was no significant difference in the maximal NPQ and there was also no difference in the NPQ relaxation dynamics (Fig. 3A). Furthermore, the OE-VDE lines show higher quantum yield of regulated energy dissipation in PSII, Y(NPQ) and lower quantum yield of nonregulated energy dissipation in PSII, Y(NO) (Fig. 3B, 3D), though we found no difference in the effective PS II quantum yield, Y(II) (Fig. 3C)[63]. Similarly, OE-VDE lines could also speed up the induction of NPQ in rice grown under high light (HL) (Fig. 3E∼3H) and in the paddy field (Fig. 3I∼3L). These results indicated that overexpression of VDE would speed up the NPQ initiation in rice exposed to excess light than WT.

**Fig 3.**
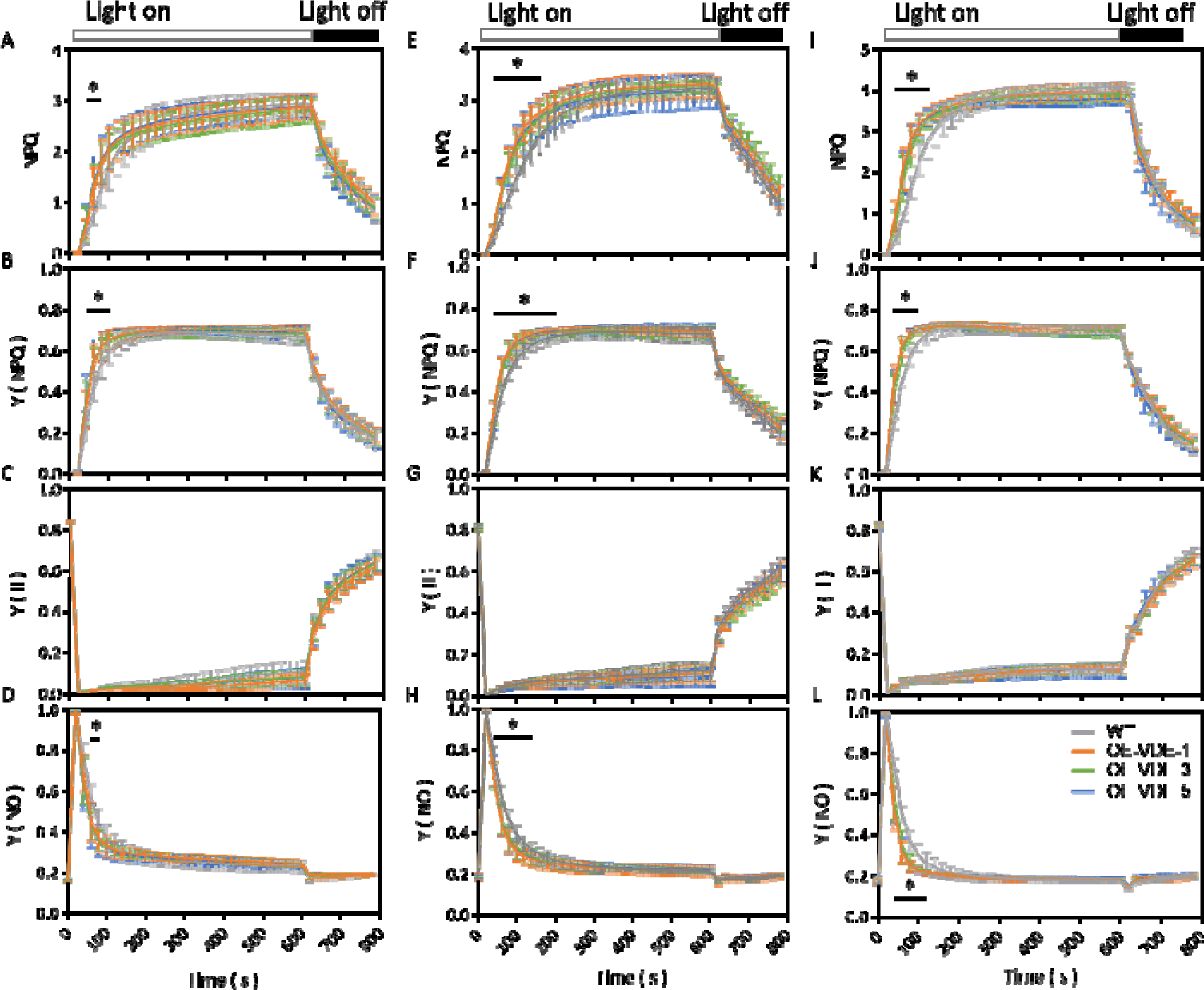
The comparison of the photosynthetic parameters based on chlorophyll fluorescence between OE-VDE lines and WT. (A∼D) Growing under normal light (400 μmol m^-2^ s^-1^, n = 10 biological replicates) all the time, (E∼H) high light (1000 μmol m^-2^ s^-1^, n = 11 biological replicates) for 3 days, (I∼L) natural light (in field, Shanghai, n = 7∼10 biological replicates). NPQ: non-photochemical quenching; Y(NPQ): the quantum yield of regulated energy dissipation in PSII; Y(II): the effective PS II quantum yield; Y(NO): the quantum yield of nonregulated energy dissipation in PSII. The fully expanded rice leaves at tillering stage were selected and went through a dark-adaption for more than 2 h before measurement. The time course of chlorophyll fluorescence was recorded every 15 s when exposed to a light of 1300 μmol m^-2^ s^-1^ for 10 min and followed by 6 min in darkness by Dual-PAM-100. *: p<0.05 (Student’s t test), the black lines represent a period of time on the x-axis.

### Overexpression of VDE accelerates photosynthesis induction from low to high light

We further examined the effect of overexpressing VDE on photosynthetic properties. We studied photosynthetic rates of rice plants grown in green house during the early tillering stage. We found no significant difference in the rates of leaf CO_2_ uptake between WT and OE-VDE lines in light response curves (*A*-Q curves) (Fig. 4A). Similarly, the NPQ and other chlorophyll fluorescence-based parameters showed no significant change (Fig. 4B∼4F). These results suggested that OE-VDE lines did not alter the photosynthetic capacity of rice under steady state light condition.

**Fig 4.**
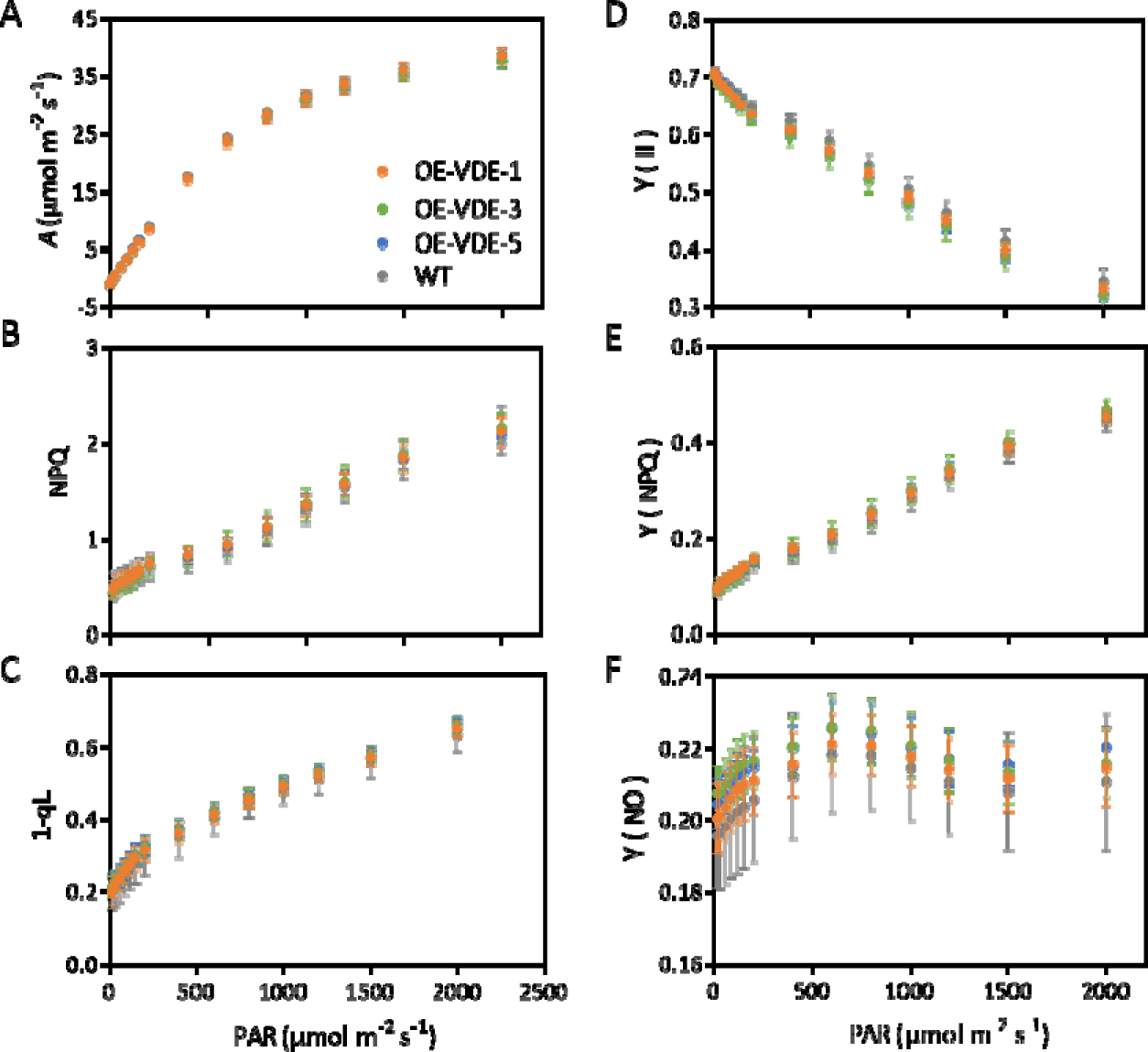
Comparation of photosynthesis parameters at stable state under different light in the OE-VDE lines and WT. (A) Net leaf CO_2_ assimilation at light saturation (A-Q); (B) NPQ, non-photochemical quenching; (C)1-qL, the redox state of Q_A_, which was estimated by Fo; (D) Y(II), the effective PS II quantum yield; (E) Y(NPQ), the quantum yield of regulated energy dissipation in PSII; (F) Y(NO), the quantum yield of nonregulated energy dissipation in PSII. The last fully expanded leaf of rice at tillering stage in the greenhouse was treated by a light of 2000 μmol m^-2^ s^-1^ for 10 min and the gas exchange and fluorescence parameters were recorded. Subsequently, the light intensity was changed to 1500, 1200, 1000, 800, 600, 400, 200, 150, 120, 90, 60, 30, 15, 0 μmol m^-2^ s^-1^ for 75 s respectively and the related gas exchange and fluorescence parameters were recorded. After that, the rice went through a dark adaptation for 2 h and the Fo and Fm were measured for calculating the related fluorescence parameters. n = 8∼9 biological replicates.

To further quantify the potential impact of altering VDE expression on NPQ and photosynthesis dynamics in the OE-VDE lines, we examined the dynamic changes of photosynthesis during the switch from low to high light, which causes a gradual increase in photosynthetic CO_2_ uptake rate [2]. The OE-VDE lines show a faster induction of photosynthesis during the switch from low light (100 μmol m^-2^ s^-1^) to high light (2000 μmol m^-2^ s^-1^). In the initial stage of transition (the 3^rd^ ∼249^th^ s), the leaf CO_2_ assimilation rate (*A*) was significantly higher in OE-VDE lines compared with WT and the percentage increase of *A* was gradually diminished along with the prolongation of rice plants exposed to high light (Fig. 5A). 250 second after the low to high light switch, we found no difference in *A* between OE-VDE lines and WT. As a result, the cumulative *A* was 20∼23% higher in the OE-VDE lines compared to WT (Fig. S3). Additionally, the g_s_ in OE-VDE lines was higher than that in WT (Fig. 5C), which might have contributed to increased CO_2_ influx [66].

**Fig 5.**
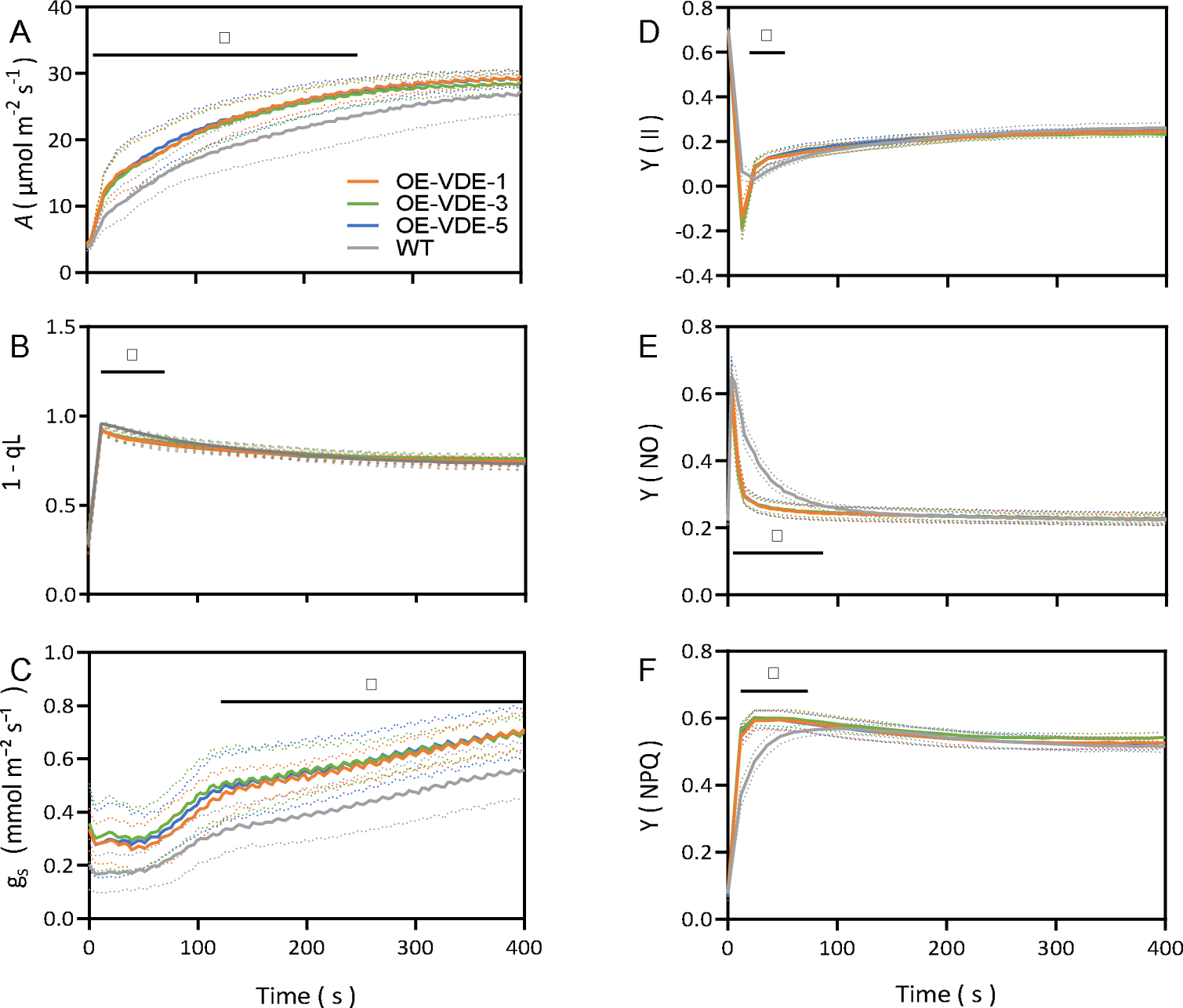
The response curves of photosynthesis and chlorophyll fluorescence parameters. (A) Dynamic net leaf CO_2_ uptake rate; (B) 1-qL, which was estimated by Fo; (C) g_s_, stomatal conductance; (D) Y(II), the effective PS II quantum yield (When the light intensity increases, the real-time fluorescence will increase at first and then decrease to a stable level, resulting that Y(II) is recorded as a negative value); (E). Y(NO), the quantum yield of nonregulated energy dissipation in PSII; (F) Y(NPQ), the quantum yield of regulated energy dissipation in PSII. The last fully expanded leaves of rice at tillering stage in the greenhouse were treated by a light of 100 μmol m^-2^ s^-1^ for 4 min, and the gas exchange and fluorescence parameters were recorded. Subsequently, the light intensity was changed to 2000 μmol m^-2^ s^-1^ for 10 min and the related gas exchange parameters were recorded every 3 s and fluorescence parameters were recorded every 12 s. After that, the rice went through a dark adaptation for more than 2 h and the Fo and Fm were recorded for calculating the related fluorescence parameters. n = 6∼8 biological replicates, *: *p*<0.05 (Student’s t test), the black lines represent a period of time on the x-axis.

### Overexpressing of VDE alleviated photoinhibition from low to high light

We further quantified the effect of OE-VDE on photoinhibition, another major factor that controls canopy photoysnthesis. Specifically, we measured chlorophyll fluorescence-based parameters during the switch process. Similar to the results of NPQ induction curve, higher Y(NPQ) and lower Y(NO) were found during photosynthesis induction in the OE-VDE lines (Fig. 5E, 5F). We also estimated 1-qL, an indicator of Q_A_ redox status [63]. The results showed that 1-qL of rice leaves in OE-VDE lines was significantly lower in the 12^th^ ∼72^nd^ s of the light switch than that in WT (Fig. 5B), which indicates that in the first 12^th^ ∼72^nd^ s the PSII center of OE-VDE lines have higher fraction of open centers [63, 67]. To gain a more accurate measurement of qL, we also used a far-red light during photosynthesis measurement. Since the difference in 1-qL was shown during the 12^th^ ∼ 72^nd^ s between the OE-VDE and WT, we chose the 33^rd^ and 336^th^ s after the light transition to give the far-red light. Results showed that the conclusions remain unchanged (Fig. S4A∼4B), i.e., the OE-VDE lines had higher DES and lower Y(NO) (Fig. 2D, 3D), which indicated the alleviated photoinhibition under high light.

### Overexpression of VDE increased biomass production in the field

Given the faster photosynthesis induction in the OE-VDE lines and increased cumulative *A* than WT during a low to high light transition, we further examined the agronomic traits of overexpression of VDE in paddy field. Compared with WT, although the plant height decreased slightly in the OE-VDE lines, the OE-VDE rice exhibited more panicles, resulting in 11∼16% increase in biomass (Fig. 6A∼6D). No significant difference in the grain yield, specific leaf weight (SLW) and leaf nitrogen content (LNC) (Fig. 6E∼6F) between WT and OE-VDE lines was observed.

**Fig 6.**
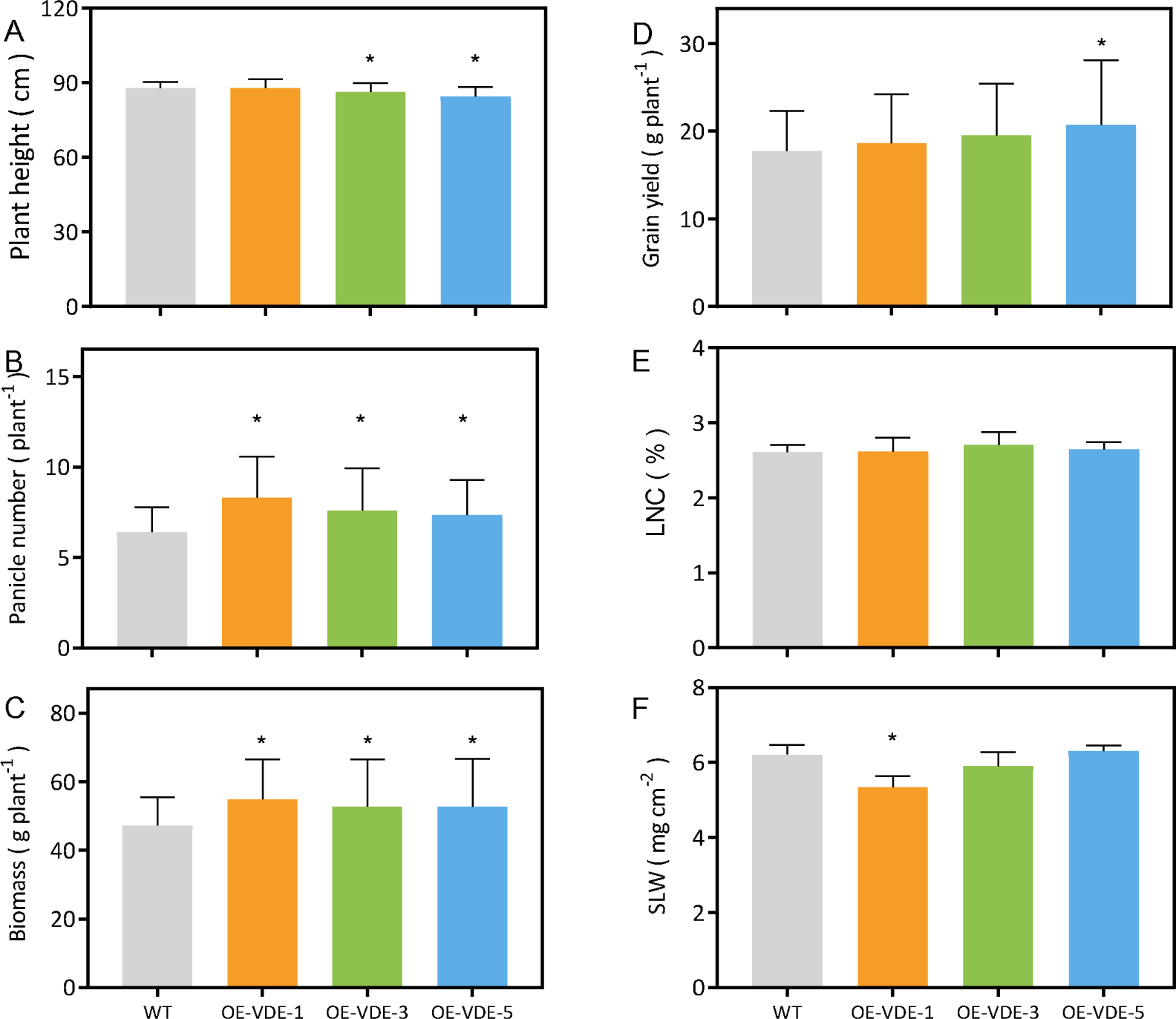
The comparison of the agronomic traits between OE-VDE lines and WT in the field. (A) Plant height (n = 40∼54), (B) panicle number (n = 40∼54), (C) biomass (n = 40∼54), (D) grain yield (n = 40∼54), (E) leaf nitrogen content (n = 5), (F) specific leaf weight (n = 5). (A∼D) The rice at harvesting stage in the field were used. (E, F) The last fully expanded leaves of rice at heading stage were used. n: biological replicates, *: *p*<0.05 (Student’s t test).

## Discussion

During the switch from low to high light, it takes time to activate many photosynthetic processes [3, 17, 21, 68, 69]. The ATP and NADPH can be rapidly increased during the initial stage of the light switch, resulting in the overly reduced PQ pool [63], while the Rubisco activation process, the NPQ dynamics and stomatal conductance dynamics all need time to adjust to new light levels [1, 2, 7, 9, 11, 13, 25, 27, 70, 71]. Identify options to manipulate the response of photosynthesis to dynamic light is regarded as a major option to improve photosynthesis for greater yields [1, 5]. In this study, we overexpressed VDE, a critical enzyme involved in the xanthophyll cycle, and found that this manipulation resulted in increased photosynthesis and biomass production (Fig. 5A, 6C). Evidences from our data suggested that the improved photosynthesis and biomass production might be due to two factors, i.e., the increased photosynthesis induction and decreased photoinhibition.

In our study, we found that acceleration of the NPQ initiation in the OE-VDE lines were accompanied by a simultaneous increase in the speed of photosynthesis induction (Fig. 5C). In the OE-VDE lines, we observed a faster response of NPQ, which resulted in more oxidized PQ pool, indicated by a decreased in 1-qL (Fig. 5B, 5C) [33, 63] and higher Y(II), which represents faster electron transport compared with WT. In line with this, we also observed higher CO_2_ assimilation during low to high light transition [65, 72-75] (Fig. 5A, 5C and 5D). In addition to the increase in Y(II), another factor that might have contributed to the faster photosynthesis induction is the increased stomatal conductance (Fig. 5C). The higher g_s_ might also be related to the lower 1-qL in the OE-VDE rice lines (Fig. 5B, C), since the redox status of PQ pool is closely linked to the stomatal conductance [76-79].

In addition to the increased speed of photosynthesis induction, the OE-VDE rice lines also show less photoinhibition. In the OE-VDE lines, the contents of Zea increased more than two times (Fig. 2C). Zea is a major factor for the initiation and relaxation of NPQ, and also a major component of qI [43]. In our OE-VDE lines, besides the faster NPQ initiation, we also found a decreased photoinhibition during a low to high light transition (Fig. 5C, 5D). The higher level of Zea and DES in the OE-VDE lines (Fig.2C, 2D) helped dissipate overloaded energy more quickly to alleviate photoinhibition. Y(NO), which represents non-regulated heat dissipation by PSII, was decreased in OE-VDE lines (Fig. 5E), which also indicates that these OE-VDE rice lines suffered less photoinhibition than WT [63, 80].

Given the high level of heterogeneity of light in a canopy [1, 2], both the decreased loss due to photoinhibition and the increased photosynthesis induction can potentially increase canopy photosynthesis and hence biomass production. Indeed, the OE-VDE lines show increased biomass production in our study (Fig. 6). This result is different from a recent report which shows that the overexpression of OsVDE in ZH11, a cultivar of rice, did not influence agronomic traits [81]. One possible reason is that the change in transcript level of VDE in the overexpression lines is different (Fig. 1B, 1C). In our study, the transcript level of VDE was increased by more than 50 times, which contrasts with the 1∼4 times increase in the earlier OE-VDE lines [81]. The different genetic background and growth conditions might also be potential reasons for the difference among agronomic traits. Though we found the different biomass production in the OE-VDE lines in our study, our OE-VDE lines also showed decreased tolerance to stress (Fig. S5), as in the earlier study [81].

In summary, this study shows that in addition to the commonly known effect of VDE on NPQ induction, it can also influence the speed of photosynthesis induction. Over-expression of VDE may represent a new approach to alter photosynthetic efficiency, by both increase the speed of its NPQ induction, which can cause alleviated photoinhibition, but also increased photosynthetic induction, both of which are preferred effect from the perspective of improving photosynthetic efficiency.

### Supplementary data

The following supplementary data are available at *JXB* online.

Fig. S1. Phylogenetic tree of VDE.

Fig. S2. Changes in the relative content of Chl and lutein in the OE-VDE lines compared with WT.

Fig. S3. The relative CO2 uptake in OE-VDE lines compared with WT from low to high light.

Fig. S4. The response of photosynthesis and 1-qL during the switch from low to high light

Fig S5. The phonotype of plants treated by 13% PEG.

Fig. S6. The result of WT and OE-VDE-1 by HPLC.

## Abbreviations

NPQ: non-photochemical quenching
LNC: leaf nitrogen content
SLW: specific leaf weight
CBB cycle: Calvin-Benson cycle
Rubisco: ribulose 1,5-bisphosphate carboxylase/oxygenase
RCA: Rubisco activase
g_m_: mesophyll conductance
g_s_: stomata conductance
s: second
h: hour
m: minute
Vio: violaxanthin
Zea: zeaxanthin
Ant: antheraxanthin
Lut: lutein
VDE: violaxanthin de-epoxidase
ZEP: Zeaxanthin epoxidase
PSII: photosystem II
LHCII: light harvested complex II
qE: energy-dependent quenching
qZ: Zea-dependent quenching
qI: photoinhibition-dependent quenching
OE-VDE-1/3/5: VDE overexpression lines
WT: wild type
HPLC: High Performance Liquid Chromatography
DES: de-epoxidation state
1-qL: the redox state of Q_A_
Y(II): the effective PS II quantum yield
Y(NPQ): the quantum yield of regulated energy dissipation in PSII
Y(NO): the quantum yield of nonregulated energy dissipation in PSII
HL: high light
*A*: CO_2_ assimilation rate
MS: Morishige & Skoog Medium.

## Acknowledgements

We acknowledged the help of Prof. Genyun Chen, Dr Qingfeng Song, Dr Tian-Gen Chang, Dr Tiangen Chang, Ms Xinyu Liu, Ms Lina Shi and other members of the plant systems biology group.

## Author contributions

XD and CF conducted experiments and wrote the paper; ZP, SZ and TQ performed experiments; ZXG conceived the study and wrote the paper.

## Acknowledgements

Authors acknowledge inspiring discussions and technical supports from Tiangen Chang, Qingfeng Song, Linxiong Mao, Fusang Liu, Genyun Chen.

---

**Fig. S1.**
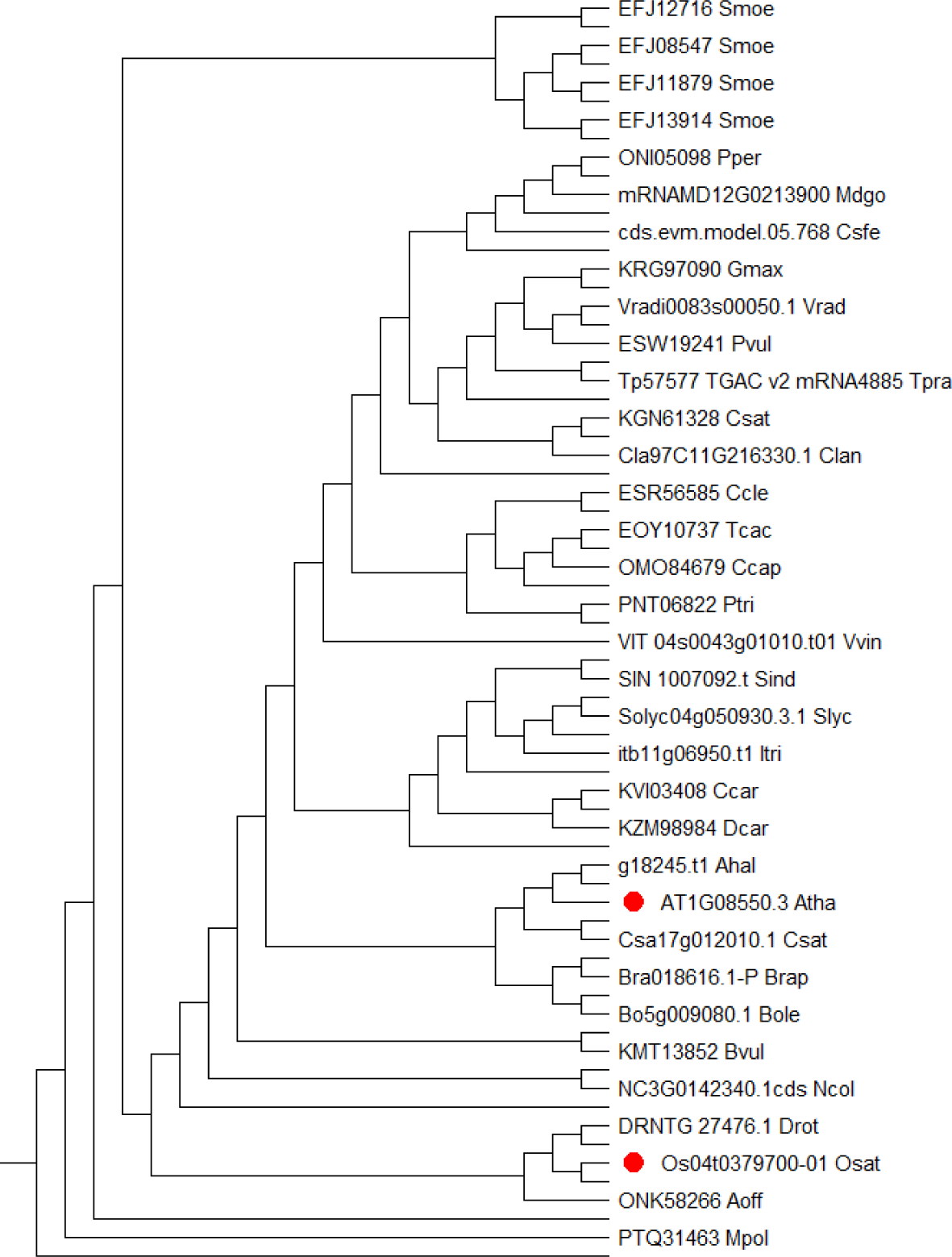
Phylogenetic tree of VDE. The data was from Ensembl Plants (http://plants.ensembl.org/index.html). The phylogenetic tree contains all of VDE’s homologues, with red circles indicating those in Arabidopsis thaliana and rice.

**Fig. S2.**
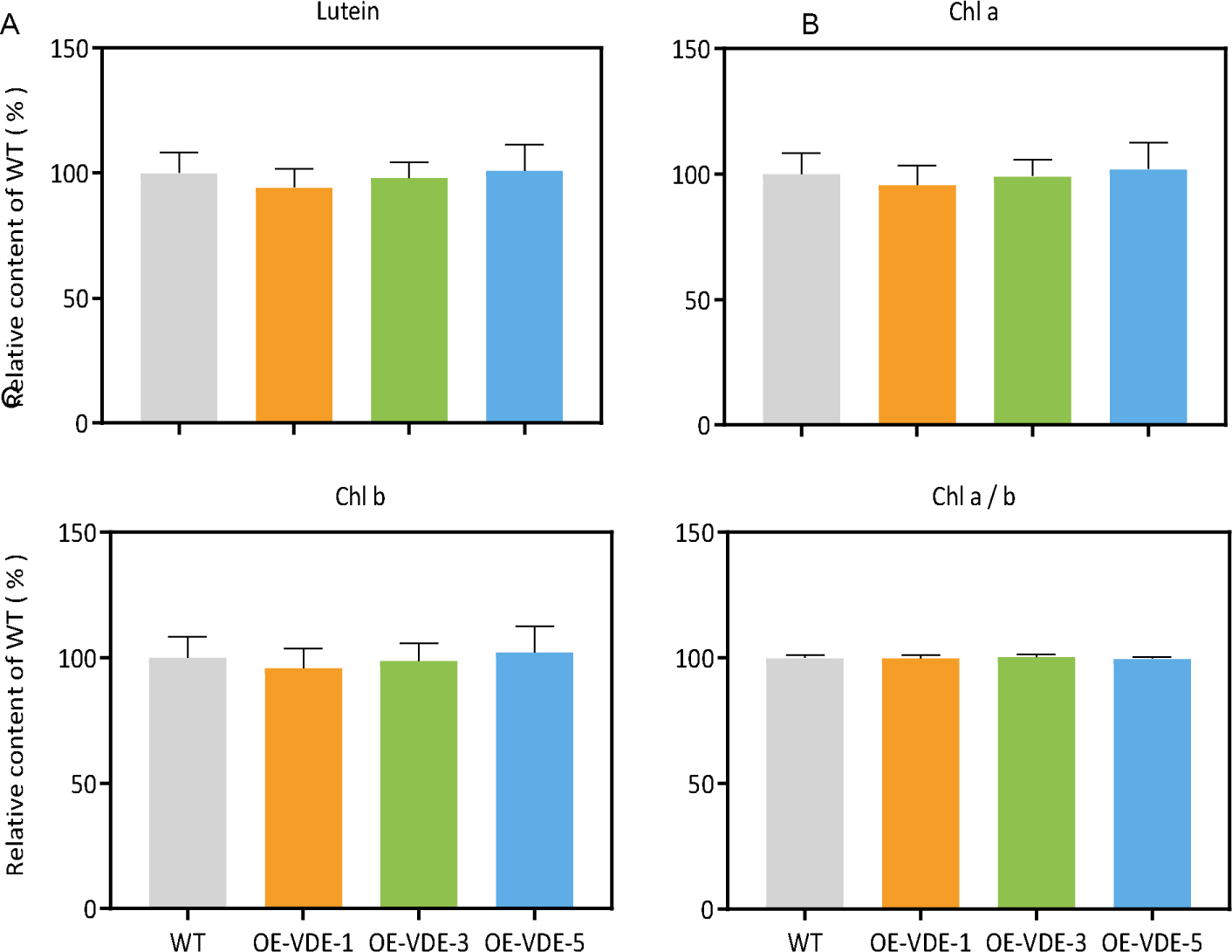
Changes in the relative content of Chl and lutein in the OE-VDE lines compared with WT. (A) Lutein, (B) Chl a, (C) Chl b, (D) Chl a/b. The last expanded rice leaves at tillering stage in the field were collected at 2 p.m. on a sunny day. n = 6∼7, *: *p*<0.05 (Student’s t test).

**Fig. S3.**
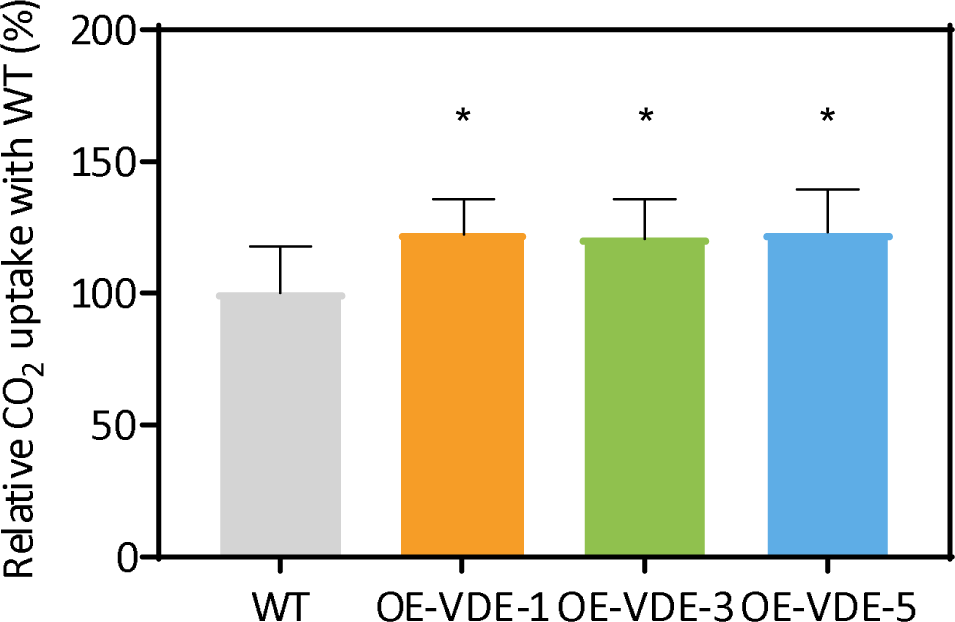
The relative CO2 uptake in OE-VDE lines compared with WT from low to high light. Calculated from the data of “Figure 3-7 A”. n = 6∼8 biological replicates, *: *p*<0.05 (Student’s t test).

**Fig. S4.**
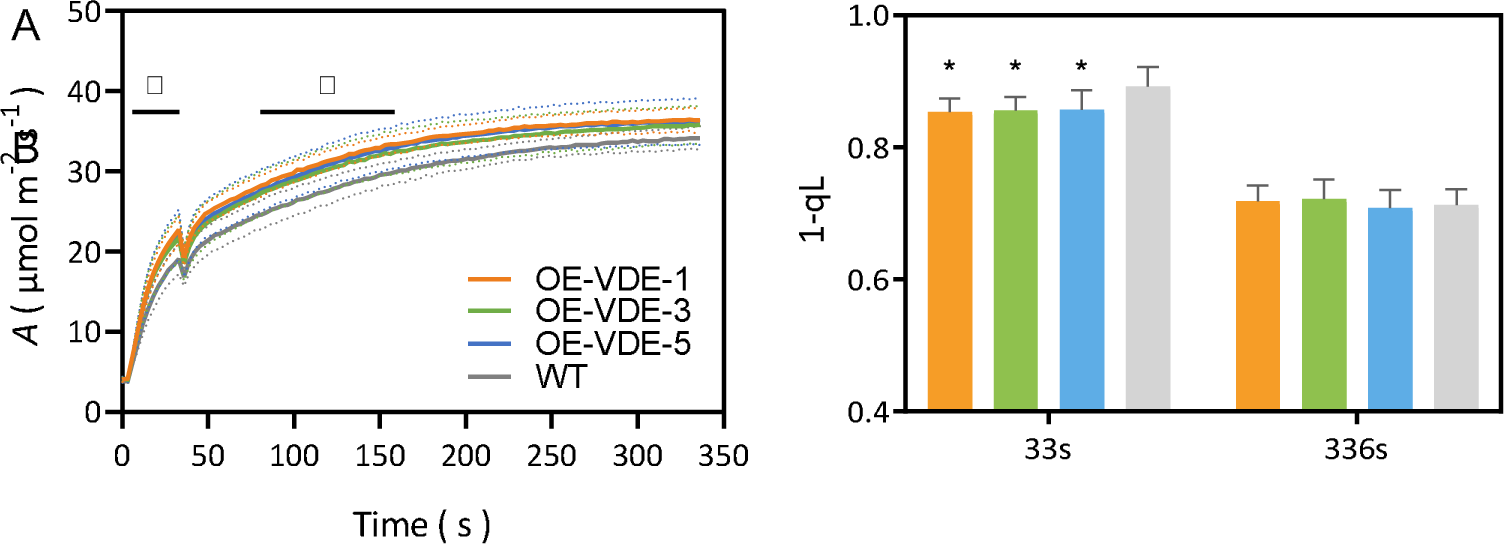
The response of photosynthesis and 1-qL during the switch from low to high light. (A) Dynamic net leaf CO2 assimilation, (B) 1-qL, calculated by Fo’. The last fully expanded leaves of rice at tillering stage in the greenhouse were treated by a light of 100 μmol m^-2^ s^-1^ for 4 min, and the gas exchange parameters were recorded. Then, the light intensity was changed to 2000 μmol m^-2^ s^-1^ for 336 s and the related gas exchange parameters were recorded every 3 s. F, Fm’ and Fo’ were recorded at 33rd s and 336th s. After that, the rice went through a dark adaptation for more than 2 h and the Fo and Fm were recorded for calculating the 1-qL. n=6∼9 biological replicates, *: *p*<0.05 (Student’s t test), the black lines represent a period of time on the x-axis.

**Fig. S5.**
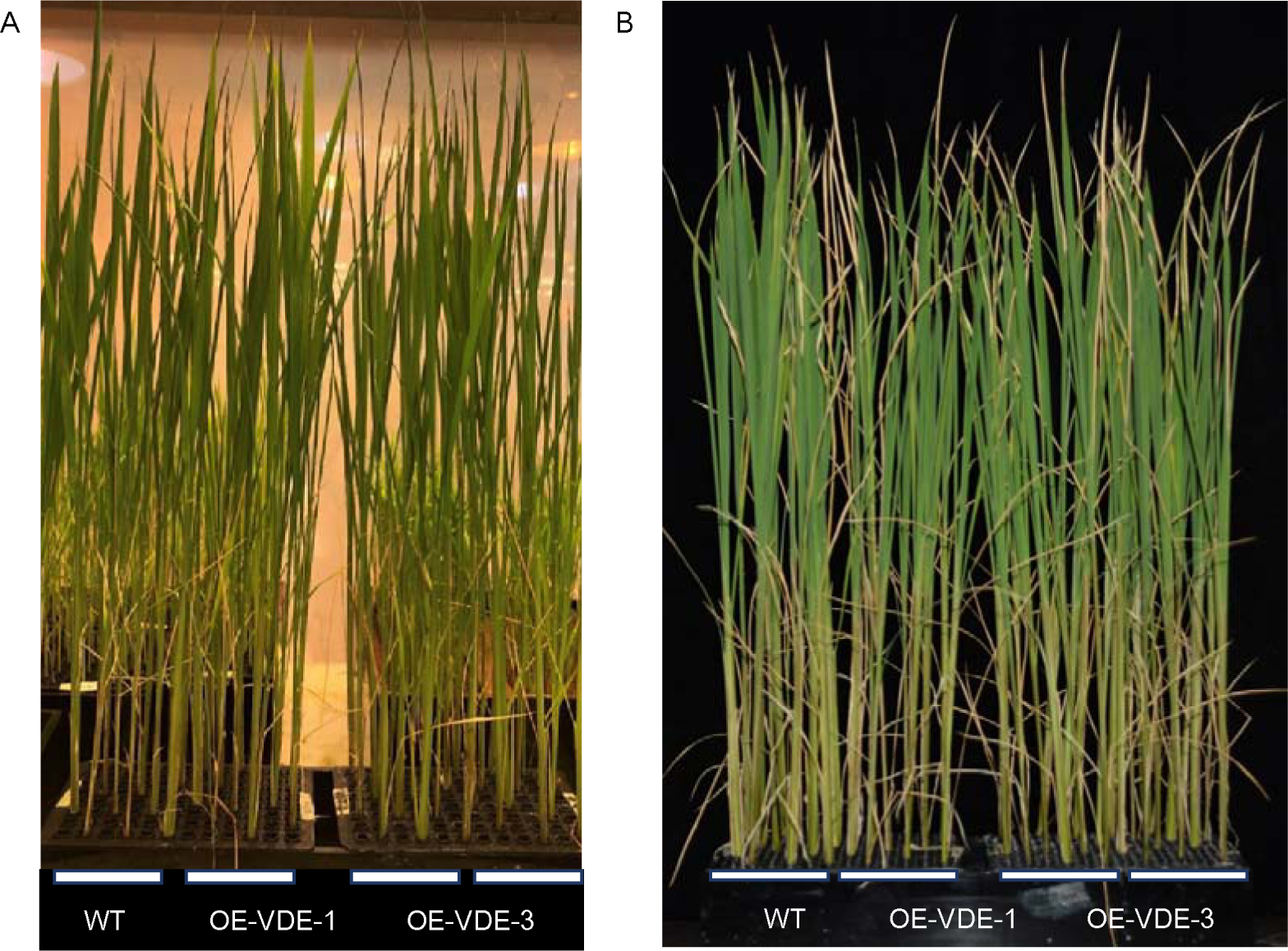
The phonotype of plants treated by 13% PEG. A. the plants before treatment. All plants were in equally high at first. B. the plants treated by 13%PEG for a week (n=12 biological replicates.). The basic liquid is ½MS.

**Fig. S6.**
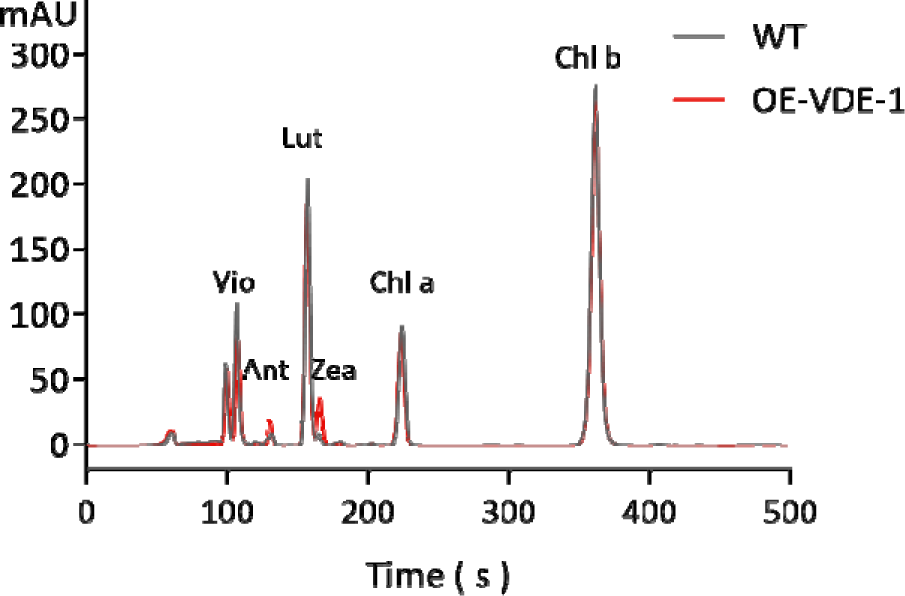
The result of WT and OE-VDE-1 by HPLC. The gray line represented the result of WT and the red line represented the result of OE-VDE-1.

## References

1. Zhu, X.G., et al., The slow reversibility of photosystem II thermal energy dissipation on transfer from high to low light may cause large losses in carbon gain by crop canopies: a theoretical analysis. J Exp Bot, 2004. 55 (400): p. 1167–75.

2. Pearcy, R.W., Sunflecks and Photosynthesis in Plant Canopies. Annual Review of Plant Physiology and Plant Molecular Biology, 1990. 41 (1): p. 421–453.

3. Yamori, W., et al., Rubisco activase is a key regulator of non-steady-state photosynthesis at any leaf temperature and, to a lesser extent, of steady-state photosynthesis at high temperature. Plant J, 2012. 71 (6): p. 871–80.

4. Soleh, M.A., et al., Identification of large variation in the photosynthetic induction response among 37 soybean [Glycine max (L.) Merr.] genotypes that is not correlated with steady-state photosynthetic capacity. Photosynth Res, 2017. 131 (3): p. 305–315.

5. Long, S.P., et al., Into the Shadows and Back into Sunlight: Photosynthesis in Fluctuating Light. Annu Rev Plant Biol, 2022. 73: p. 617–648.

6. Taylor, S.H., et al., Faster than expected Rubisco deactivation in shade reduces cowpea photosynthetic potential in variable light conditions. Nat Plants, 2022. 8(2): p. 118–124.

7. Taylor, S.H. and S.P. Long, Slow induction of photosynthesis on shade to sun transitions in wheat may cost at least 21% of productivity. Philos Trans R Soc Lond B Biol Sci, 2017. 372 (1730).

8. Yoon, D.-K., et al., Transgenic rice overproducing Rubisco exhibits increased yields with improved nitrogen-use efficiency in an experimental paddy field. Nature Food, 2020. 1(2): p. 134–139.

9. Wang, Y., et al., Photosynthesis in the fleeting shadows: an overlooked opportunity for increasing crop productivity? Plant J, 2020. 101(4): p. 874–884.

10. De Souza, A.P., et al., Photosynthesis across African cassava germplasm is limited by Rubisco and mesophyll conductance at steady state, but by stomatal conductance in fluctuating light. New Phytol, 2020. 225(6): p. 2498–2512.

11. Salter, W.T., et al., Rate of photosynthetic induction in fluctuating light varies widely among genotypes of wheat. J Exp Bot, 2019. 70(10): p. 2787–2796.

12. Deans, R.M., et al., Plant water-use strategy mediates stomatal effects on the light induction of photosynthesis. New Phytol, 2019. 222(1): p. 382–395.

13. Morales, A., et al., Dynamic modelling of limitations on improving leaf CO2 assimilation under fluctuating irradiance. Plant Cell Environ, 2018. 41(3): p. 589–604.

14. Burgess, A.J., et al., High-Resolution Three-Dimensional Structural Data Quantify the Impact of Photoinhibition on Long-Term Carbon Gain in Wheat Canopies in the Field. Plant Physiol, 2015. 169(2): p. 1192–204.

15. Sakoda, K., et al., Stomatal, mesophyll conductance, and biochemical limitations to photosynthesis during induction. Plant Physiol, 2021. 185(1): p. 146–160.

16. Yamori, W., et al., Increased stomatal conductance induces rapid changes to photosynthetic rate in response to naturally fluctuating light conditions in rice. Plant Cell Environ, 2020. 43(5): p. 1230–1240.

17. Qu, Y., et al., Overexpression of both Rubisco and Rubisco activase rescues rice photosynthesis and biomass under heat stress. Plant Cell Environ, 2021. 44(7): p. 2308–2320.

18. Deans, R.M., G.D. Farquhar, and F.A. Busch, Estimating stomatal and biochemical limitations during photosynthetic induction. Plant Cell Environ, 2019. 42(12): p. 3227–3240.

19. Mott, K.A. and I.E. Woodrow, Modelling the role of Rubisco activase in limiting non-steady-state photosynthesis. J Exp Bot, 2000. 51 Spec No: p. 399-406.

20. Portis, A.R., M.E. Salvucci, and W.L. Ogren, Activation of Ribulosebisphosphate Carboxylase/Oxygenase at Physiological CO(2) and Ribulosebisphosphate Concentrations by Rubisco Activase. Plant Physiol, 1986. 82(4): p. 967–71.

21. Yamori, W. and S. von Caemmerer, Effect of Rubisco activase deficiency on the temperature response of CO2 assimilation rate and Rubisco activation state: insights from transgenic tobacco with reduced amounts of Rubisco activase. Plant Physiol, 2009. 151(4): p. 2073–82.

22. Pons, T.L., et al., Estimating mesophyll conductance to CO_2_: methodology, potential errors, and recommendations. J Exp Bot, 2009. 60(8): p. 2217–34.

23. Kimura, H., et al., Improved stomatal opening enhances photosynthetic rate and biomass production in fluctuating light. J Exp Bot, 2020. 71(7): p. 2339–2350.

24. Hubbart, S., et al., Enhanced thylakoid photoprotection can increase yield and canopy radiation use efficiency in rice. Commun Biol, 2018. 1: p. 22.

25. Kromdijk, J., et al., Improving photosynthesis and crop productivity by accelerating recovery from photoprotection. Science, 2016. 354(6314): p. 857-861.

26. Murchie, E.H., A. Ali, and T. Herman, Photoprotection as a Trait for Rice Yield Improvement: Status and Prospects. Rice (N Y), 2015. 8(1): p. 31.

27. De Souza, A.P., et al., Soybean photosynthesis and crop yield are improved by accelerating recovery from photoprotection. Science, 2022. 377(6608): p. 851-854.

28. Horton, P. and A. Ruban, Molecular design of the photosystem II light-harvesting antenna: photosynthesis and photoprotection. J Exp Bot, 2005. 56(411): p. 365–73.

29. Jahns, P. and A.R. Holzwarth, The role of the xanthophyll cycle and of lutein in photoprotection of photosystem II. Biochim Biophys Acta, 2012. 1817(1): p. 182–93.

30. Niyogi, K.K. and T.B. Truong, Evolution of flexible non-photochemical quenching mechanisms that regulate light harvesting in oxygenic photosynthesis. Curr Opin Plant Biol, 2013. 16(3): p. 307–14.

31. Ruban, A.V., Nonphotochemical Chlorophyll Fluorescence Quenching: Mechanism and Effectiveness in Protecting Plants from Photodamage. Plant Physiol, 2016. 170(4): p. 1903–16.

32. Armbruster, U., et al., Ion antiport accelerates photosynthetic acclimation in fluctuating light environments. Nat Commun, 2014. 5: p. 5439.

33. Finazzi, G., et al., Ions channels/transporters and chloroplast regulation. Cell Calcium, 2015. 58(1): p. 86–97.

34. Demmig-Adams, B. and W.W. Adams, The role of xanthophyll cycle carotenoids in the protection of photosynthesis. Trends in Plant Science,> 1996. 1(1): p. 21–26.

35. Li, X.P., et al., A pigment-binding protein essential for regulation of photosynthetic light harvesting. Nature, 2000. 403(6768): p. 391-395.

36. Rockholm, D.C. and H.Y. Yamamoto, Violaxanthin De-Epoxidase (Purification of a 43-Kilodalton Lumenal Protein from Lettuce by Lipid-Affinity Precipitation with Monogalactosyldiacylglyceride). Plant Physiology, 1996. 110(2): p. 697–703.

37. Goss, R. and B. Lepetit, Biodiversity of NPQ. J Plant Physiol, 2015. 172: p. 13–32.

38. Büch, K., H. Stransky, and A. Hager, FAD is a further essential cofactor of the NAD(P)H and O2-dependent zeaxanthin-epoxidase. FEBS Letters, 1995. 376(1-2): p. 45–48.

39. Li, X.P., et al., PsbS-dependent enhancement of feedback de-excitation protects photosystem II from photoinhibition. Proc Natl Acad Sci U S A, 2002. 99(23): p. 15222–7.

40. Li, X.P., et al., Regulation of photosynthetic light harvesting involves intrathylakoid lumen pH sensing by the PsbS protein. J Biol Chem, 2004. 279(22): p. 22866–74.

41. Horton, P., M. Wentworth, and A. Ruban, Control of the light harvesting function of chloroplast membranes: the LHCII-aggregation model for non-photochemical quenching. FEBS Lett, 2005. 579(20): p. 4201–6.

42. Dong, L., et al., The PsbS protein plays important roles in photosystem II supercomplex remodeling under elevated light conditions. J Plant Physiol, 2015. 172: p. 33–41.

43. Demmig-Adams, B., et al., Zeaxanthin, a Molecule for Photoprotection in Many Different Environments. Molecules, 2020. 25(24).

44. Nawrocki, W.J., et al., Molecular origins of induction and loss of photoinhibition-related energy dissipation qI. Sci Adv, 2021. 7(52): p. eabj0055.

45. Murchie, E.H. and K.K. Niyogi, Manipulation of photoprotection to improve plant photosynthesis. Plant Physiol, 2011. 155(1): p. 86–92.

46. Murata, N. and Y. Nishiyama, ATP is a driving force in the repair of photosystem II during photoinhibition. Plant Cell Environ, 2018. 41(2): p. 285–299.

47. Tyystjarvi, E., Photoinhibition of Photosystem II. Int Rev Cell Mol Biol, 2013. 300: p. 243–303.

48. Kok, B., On the inhibition of photosynthesis by intense light. Biochim Biophys Acta, 1956. 21(2): p. 234–44.

49. Aro, E.M., I. Virgin, and B. Andersson, Photoinhibition of Photosystem II. Inactivation, protein damage and turnover. Biochim Biophys Acta, 1993. 1143(2): p. 113–34.

50. Krah, N.M. and B.A. Logan, Loss of psbS expression reduces vegetative growth, reproductive output, and light-limited, but not light-saturated, photosynthesis in Arabidopsis thaliana (Brassicaceae) grown in temperate light environments. Am J Bot, 2010. 97(4): p. 644–9.

51. Gao, S., et al., Overexpression and suppression of violaxanthin de-epoxidase affects the sensitivity of photosystem II photoinhibition to high light and chilling stress in transgenic tobacco. J Integr Plant Biol, 2010. 52(3): p. 332–9.

52. Cao, Y., et al., Overexpression of zeaxanthin epoxidase gene from Medicago sativa enhances the tolerance to low light in transgenic tobacco. Acta Biochim Pol, 2018. 65(3): p. 431–435.

53. Garcia-Molina, A. and D. Leister, Accelerated relaxation of photoprotection impairs biomass accumulation in Arabidopsis. Nat Plants, 2020. 6(1): p. 9–12.

54. Lehretz, G.G., et al., High non-photochemical quenching of VPZ transgenic potato plants limits CO2 assimilation under high light conditions and reduces tuber yield under fluctuating light. J Integr Plant Biol, 2022.

55. Guan, C., et al., Positive feedback regulation of a Lycium chinense-derived VDE gene by drought-induced endogenous ABA, and over-expression of this VDE gene improve drought-induced photo-damage in Arabidopsis. J Plant Physiol, 2015. 175: p. 26–36.

56. Li, X., et al., Molecular cloning and characterization of violaxanthin de-epoxidase (CsVDE) in cucumber. PLoS One, 2013. 8(5): p. e64383.

57. Sun, L.N., et al., Overexpression of the ChVDE gene, encoding a violaxanthin de-epoxidase, improves tolerance to drought and salt stress in transgenic Arabidopsis. 3 Biotech, 2019. 9(5): p. 197.

58. Yang, S., et al., Peanut violaxanthin de-epoxidase alleviates the sensitivity of PSII photoinhibition to heat and high irradiance stress in transgenic tobacco. Plant Cell Rep, 2015. 34(8): p. 1417–28.

59. Hallin, E.I., K. Guo, and H.E. Akerlund, Violaxanthin de-epoxidase disulphides and their role in activity and thermal stability. Photosynth Res, 2015. 124(2): p. 191–8.

60. Chen, Z. and D.R. Gallie, Violaxanthin de-epoxidase is rate-limiting for non-photochemical quenching under subsaturating light or during chilling in Arabidopsis. Plant Physiol Biochem, 2012. 58: p. 66–82.

61. Hiei, Y., et al., Efficient transformation of rice (Oryza sativa L.) mediated by Agrobacterium and sequence analysis of the boundaries of the T-DNA. Plant J, 1994. 6(2): p. 271–82.

62. Yudina, L., et al., A light-induced decrease in the photochemical reflectance index (PRI) can be used to estimate the energy-dependent component of non-photochemical quenching under heat stress and soil drought in pea, wheat, and pumpkin. Photosynth Res, 2020. 146(1-3): p. 175–187.

63. Kramer, D.M., et al., New Fluorescence Parameters for the Determination of QA Redox State and Excitation Energy Fluxes. Photosynth Res, 2004. 79(2): p. 209.

64. Hendrickson, L., R.T. Furbank, and W.S. Chow, A simple alternative approach to assessing the fate of absorbed light energy using chlorophyll fluorescence. Photosynth Res, 2004. 82(1): p. 73–81.

65. Demmig-Adams, B. and W.W. Adams, Xanthophyll cycle and light stress in nature: uniform response to excess direct sunlight among higher plant species. Planta, 1996. 198(3): p. 460–470.

66. Messinger, S.M., T.N. Buckley, and K.A. Mott, Evidence for involvement of photosynthetic processes in the stomatal response to CO2. Plant Physiol, 2006. 140(2): p. 771–8.

67. Oxborough, K. and N.R. Baker, Resolving chlorophyll a fluorescence images of photosynthetic efficiency into photochemical and non-photochemical components – calculation of qP and Fv/Fm without measuring Fo. Photosynthesis Research, 1997. 54(2): p. 135–142.

68. Iniguez, C., P. Aguilo-Nicolau, and J. Galmes, Improving photosynthesis through the enhancement of Rubisco carboxylation capacity. Biochem Soc Trans, 2021. 49(5): p. 2007–2019.

69. Kaiser, E., et al., Dynamic photosynthesis in different environmental conditions. J Exp Bot, 2015. 66(9): p. 2415–26.

70. Bailey-Serres, J., et al., Genetic strategies for improving crop yields. Nature, 2019. 575(7781): p. 109–118.

71. Acevedo-Siaca, L.G., et al., Variation between rice accessions in photosynthetic induction in flag leaves and underlying mechanisms. J Exp Bot, 2021. 72(4): p. 1282–1294.

72. Havaux, M., Plastoquinone In and Beyond Photosynthesis. Trends Plant Sci, 2020. 25(12): p. 1252–1265.

73. Borisova-Mubarakshina, M.M., D.V. Vetoshkina, and B.N. Ivanov, Antioxidant and signaling functions of the plastoquinone pool in higher plants. Physiol Plant, 2019. 166(1): p. 181–198.

74. Murchie, E.H. and A.V. Ruban, Dynamic non-photochemical quenching in plants: from molecular mechanism to productivity. Plant J,> 2020. 101(4): p. 885–896.

75. Miyake, C., et al., Acclimation of tobacco leaves to high light intensity drives the plastoquinone oxidation system--relationship among the fraction of open PSII centers, non-photochemical quenching of Chl fluorescence and the maximum quantum yield of PSII in the dark. Plant Cell Physiol, 2009. 50(4): p. 730–43.

76. Glowacka, K., et al., Photosystem II Subunit S overexpression increases the efficiency of water use in a field-grown crop. Nat Commun, 2018. 9(1): p. 868.

77. Dwyer, S.A., et al., Antisense reductions in the PsbO protein of photosystem II leads to decreased quantum yield but similar maximal photosynthetic rates. J Exp Bot, 2012. 63(13): p. 4781–95.

78. von Caemmerer, S., et al., Stomatal conductance does not correlate with photosynthetic capacity in transgenic tobacco with reduced amounts of Rubisco. J Exp Bot, 2004. 55(400): p. 1157–66.

79. Baroli, I., et al., The contribution of photosynthesis to the red light response of stomatal conductance. Plant Physiol, 2008. 146(2): p. 737–47.

80. Baker, N.R., Chlorophyll fluorescence: a probe of photosynthesis in vivo. Annu Rev Plant Biol, 2008. 59: p. 89–113.

81. Wang, X., et al., OsVDE, a xanthophyll cycle key enzyme, mediates abscisic acid biosynthesis and negatively regulates salinity tolerance in rice. Planta, 2021. 255(1): p. 6.

